# A streamlined method for transposon mutagenesis of *Rickettsia parkeri* yields numerous mutations that impact infection

**DOI:** 10.1101/277160

**Authors:** Rebecca L. Lamason, Natasha M. Kafai, Matthew D. Welch

## Abstract

The rickettsiae are obligate intracellular alphaproteobacteria that exhibit a complex infectious life cycle in both arthropod and mammalian hosts. As obligate intracellular bacteria, *Rickettsia* are highly adapted to living inside a variety of host cells, including vascular endothelial cells during mammalian infection. Although it is assumed that the rickettsiae produce numerous virulence factors that usurp or disrupt various host cell pathways, they have been challenging to genetically manipulate to identify the key bacterial factors that contribute to infection. Motivated to overcome this challenge, we sought to expand the repertoire of available rickettsial loss-of-function mutants, using an improved *mariner*-based transposon mutagenesis scheme. Here, we present the isolation of over 100 transposon mutants in the spotted fever group species *Rickettsia parkeri*. These mutants targeted genes implicated in a variety of pathways, including bacterial replication and metabolism, hypothetical proteins, the type IV secretion system, as well as factors with previously established roles in host cell interactions and pathogenesis. Given the need to identify critical virulence factors, forward genetic screens such as this will provide an excellent platform to more directly investigate rickettsial biology and pathogenesis.

## Introduction

The *Rickettsia* are a genus of obligate intracellular alphaproteobacteria that are divided into four groups - the spotted fever group (SFG), typhus group (TG), ancestral group (AG), and transitional group (TRG) [1]. They inhabit arthropods (ticks, fleas, and mites), and many can be transmitted to humans and other mammals. Pathogenic species primarily target endothelial cells in the vasculature, causing a variety of vascular diseases such as typhus and Rocky Mountain spotted fever [2]. Despite the prevalence of rickettsial diseases throughout the world, we know little about the bacterial factors required for infection and pathogenesis.

The SFG species *Rickettsia parkeri,* a tick-borne pathogen that causes a mild form of spotted fever in humans [3,4], is emerging as a model organism to study SFG rickettsial pathogenesis. *R. parkeri* can be studied under BL2 conditions and has animal models of pathogenesis that mimic aspects of human infection [5,6]. Furthermore, the *R. parkeri* life cycle closely matches that of the more virulent SFG species *R. rickettsii* [7,8], the causative agent of Rocky Mountain spotted fever. Like its more virulent relative, *R. parkeri* invades non-phagocytic cells and is taken into a primary phagocytic vacuole [9]. They then break out of this vacuole and enter the cytosol to replicate and grow [10]. *R. parkeri* and many other *Rickettsia* species also hijack the host cell actin cytoskeleton to polymerize actin tails and undergo actin-based motility [11-13]. During spread, motile *R. parkeri* move to a host cell-cell junction but then lose their actin tails before entering into a short (∼1 µm) plasma membrane protrusion that is subsequently engulfed into the neighboring cell. The bacterium then lyses the double-membrane secondary vacuole to enter the neighboring cell cytosol and begin the process again [14]. Because of its experimental tractability and the fact that its lifecycle is indistinguishable from more virulent species, *R. parkeri* provides an attractive system for investigating rickettsial host-pathogen interactions.

As an obligate intracellular pathogen, *R. parkeri* must produce virulence factors that usurp or disrupt various host cell pathways. However, due to their obligate growth requirement, it has been challenging to genetically manipulate the rickettsiae to identify the key bacterial factors that contribute to infection [15]. Fortunately, recent advances have expanded the genetic toolkit that can be used in the rickettsiae, allowing us to more directly peer into the molecular mechanisms that drive rickettsial biology. Chief among these advances was the development of a *Himar1 mariner*-based transposon system for random mutagenesis of rickettsial genomes [16]. To date, smaller-scale mutagenesis studies have been completed in the TG species *R. prowazekii* [16-18] and the SFG species *R. rickettsii* [18,19].

Despite these advances, we still do not know all of the critical bacterial factors that mediate interactions with the host. Moreover, many of the genes in *R. parkeri* are annotated to encode hypothetical proteins, which limits our ability to rationally explore their functions. Therefore, we set out to expand the repertoire of available *R. parkeri* mutants using a forward genetic screen. We used the *mariner*-based transposon system [16] and developed a more streamlined protocol to rapidly isolate *R. parkeri* mutants that alter plaque size. To date, we have isolated over 100 mutants that disrupt genes predicted to function in a variety of pathways. We have previously published our detailed analysis of three mutants – in *sca2, rickA,* and *sca4* [14,20]. Here, we present the full panel of mutants to demonstrate the potential and ease of developing rickettsial transposon libraries.

## Materials and methods

### Cell lines

Vero cells (monkey, kidney epithelial) were obtained from the University of California, Berkeley tissue culture facility and grown in Dulbecco’s modified Eagle’s medium (DMEM) (Invitrogen) containing 5% fetal bovine serum (FBS) at 37°C in 5% CO_2_.

### Transposon mutagenesis in *R. parkeri*

*R. parkeri* Portsmouth strain was a gift from Dr. Chris Paddock (Centers for Disease Control and Prevention). Wild-type *R. parkeri* were expanded and purified by centrifugation through a 30% MD-76R solution, as previously described [14]. The pMW1650 plasmid carrying the *Himar1 mariner*-based transposon [16] (a gift from Dr. David Wood, University of South Alabama) was used to generate *R. parkeri* strains carrying transposon insertions. To isolate small plaque mutants, a small-scale electroporation protocol was designed. First, a T75 cm^2^ flask of confluent Vero cells was infected with approximately 10^7^ WT *R. parkeri*. Three days post-infection, when Vero cells were at least 90% rounded, they were scraped from the flask. Infected cells were spun down for 5 min at 1800 x g at 4°C and resuspended in 3-6 ml K-36 buffer. To mechanically disrupt infected cells and release bacteria, cells were either passed through a 27.5 gauge syringe needle 10 times, or vortexed at ∼2900 rpm using a Vortex Genie 2 (Scientific Industries Inc.) in a 15 ml conical tube containing 2 g of 1 mm glass beads, with two 30 s pulses and 30 s incubations in ice after each pulse. This bead disruption procedure was adopted for a majority of the screen because it was faster and reduced the possibility of a needle stick. Host cell debris was pelleted for 5 min at 200 x g at 4°C. The supernatant containing *R. parkeri* was transferred to 1.5 ml microcentrifuge tubes and spun for 2 min at 9000 x g at 4°C. The bacterial pellets were then washed three times in cold 250 mM sucrose, resuspended in 50 µl cold 250 mM sucrose, mixed with 1 µg of pMW1650 plasmid, placed in a 0.1 cm cuvette, and electroporated at 1.8 kV, 200 ohms, 25 µF, 5 ms using a Gene Pulser Xcell (Bio-Rad). Bacteria were immediately recovered in 1.2 ml brain heart infusion (BHI) media. To infect confluent Vero cells in 6-well plates, media was removed from each well, and cells were washed with phosphate-buffered saline (PBS). 100 µl of electroporated bacteria was added per well, and plates were placed in a humidified chamber and rocked for 30 min at 37°C. An overlay of DMEM with 5% FBS and 0.5% agarose was added to each well. Infected cells were incubated at 33°C, 5% CO_2_ for 24 h at which point a second overlay was added containing rifampicin (Sigma) to a final concentration 200 ng/ml to select for transformants. After at least 3 or 4 d, plaques were visible by eye in the cell monolayer, and plaques smaller or bigger relative to neighboring plaques were selected for further analysis.

To isolate and amplify mutant strains and map the sites of transposon insertion, plaques were picked and resuspended in 200 µl of BHI. Media was aspirated from confluent Vero cells in 6-well plates, and the isolated plaque resuspension was used to infect the cells at 37°C for 30 min with rocking. Then 3 ml DMEM with 2% FBS and 200 ng/ml rifampicin was added to each well, and infections progressed until monolayers were fully infected. Infected cells were isolated as described above using mechanical disruption, except that bacteria were immediately resuspended in BHI without a sucrose wash and stored at -80°C.

## Semi-random nested PCR

To map the transposon insertion sites, plaque-purified *R. parkeri* strains were boiled for 10 min and used as templates for PCR reactions. Genomic DNA at insertion sites was amplified for sequencing using semi-random nested PCR. The first “external” PCR reaction used transposon-specific primers (ExTn1 5‘- CACCAATTGCTAAATTAGCTTTAGTTCC-3‘; or ExTn2 5‘-GTGAGCTATGAGAAAGCGCCACGC-3‘) and a universal primer (Univ1 5‘- GCTAGCGGCCGCACTAGTCGANNNNNNNNNNCTTCT-3‘). This yielded the“external” PCR product that served as a template in the subsequent “internal” PCR reaction using transposon-specific primers (InTn1 5‘- GCTAGCGGCCGCGGTCCTTGTACTTGTTTATAATTATCATGAG-3‘; or InTn2 5‘- GCTAGCGGCCGCCCTGGTATCTTTATAGTCCTGTCGG-3‘) and a different universal primer (Univ2 5‘-GCTAGCGGCCGCACTAGTCGA-3‘). PCR products were cleaned using ExoSAP-IT PCR Product Cleanup Reagent (Affymetrix) and sequenced using primers SR095 5‘-CGCCACCTCTGACTTGAGCGTCG-3’ and SR096 5‘-CCATATGAAAACACTCCAAAAAAC-3‘. Genomic locations were determined using BLAST against the *R. parkeri* strain Portsmouth genome (GenBank/NCBI accession NC_017044.1).

## Results

### Design of an improved transposon mutagenesis scheme

We used the pMW1650 plasmid, which carries a *Himar1 mariner*-based transposon [16], to randomly mutate the *R. parkeri* genome. pMW1650 encodes the *Himar1* transposase, a transposon cassette that contains the *R. prowazekii arr-2* rifampin resistance gene, and a variant of green fluorescent protein (GFPuv) [16] (Fig 1A). The first reported application of this system in *R. prowazekii* [16] and *R. rickettsii* [18] yielded some transposon mutants, but we sought to improve the mutagenesis scheme to increase the chances of identifying genes important for infection. Therefore, we developed a simple and rapid procedure to extract bacteria from infected host cells. In the past, we had purified *R. parkeri* from infected host cells using an hours-long process involving mechanical disruption and density gradient centrifugation prior to electroporation [21]. We designed a new procedure to isolate and electroporate bacteria and re-infect host cells in under an hour. To mechanically disrupt infected cells, we either passed infected cells through syringe needle or vortexed cells in the presence of 1 mm glass beads. Samples were then spun at low speed for 5 min to pellet host cell debris, followed by a 2 min high-speed spin to pellet bacteria. *Rickettsia* were then quickly washed 2-3 times in cold sucrose prior to electroporation.

**Fig 1.**
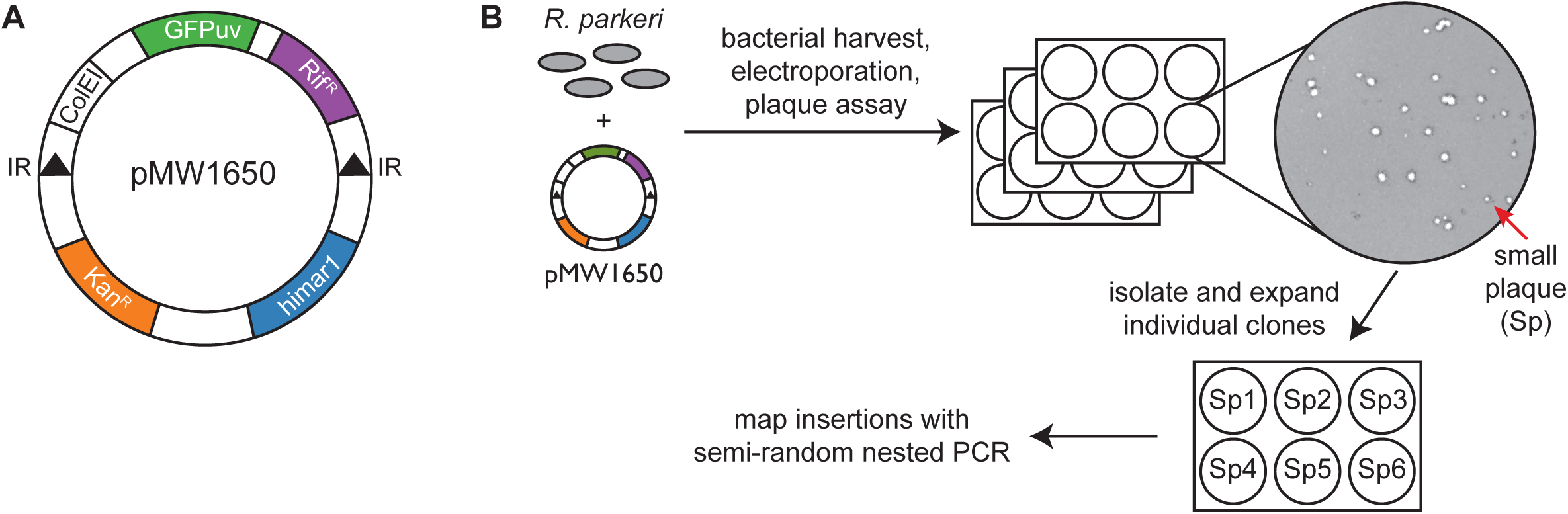
Transposon mutagenesis of *R. parkeri*. (A) Map of the pMW1650 plasmid used in this study for transposon mutagenesis (IR, inverted repeats). (B) Experimental scheme for transposon mutagenesis and isolation of individual mutants.

**Fig 2.**
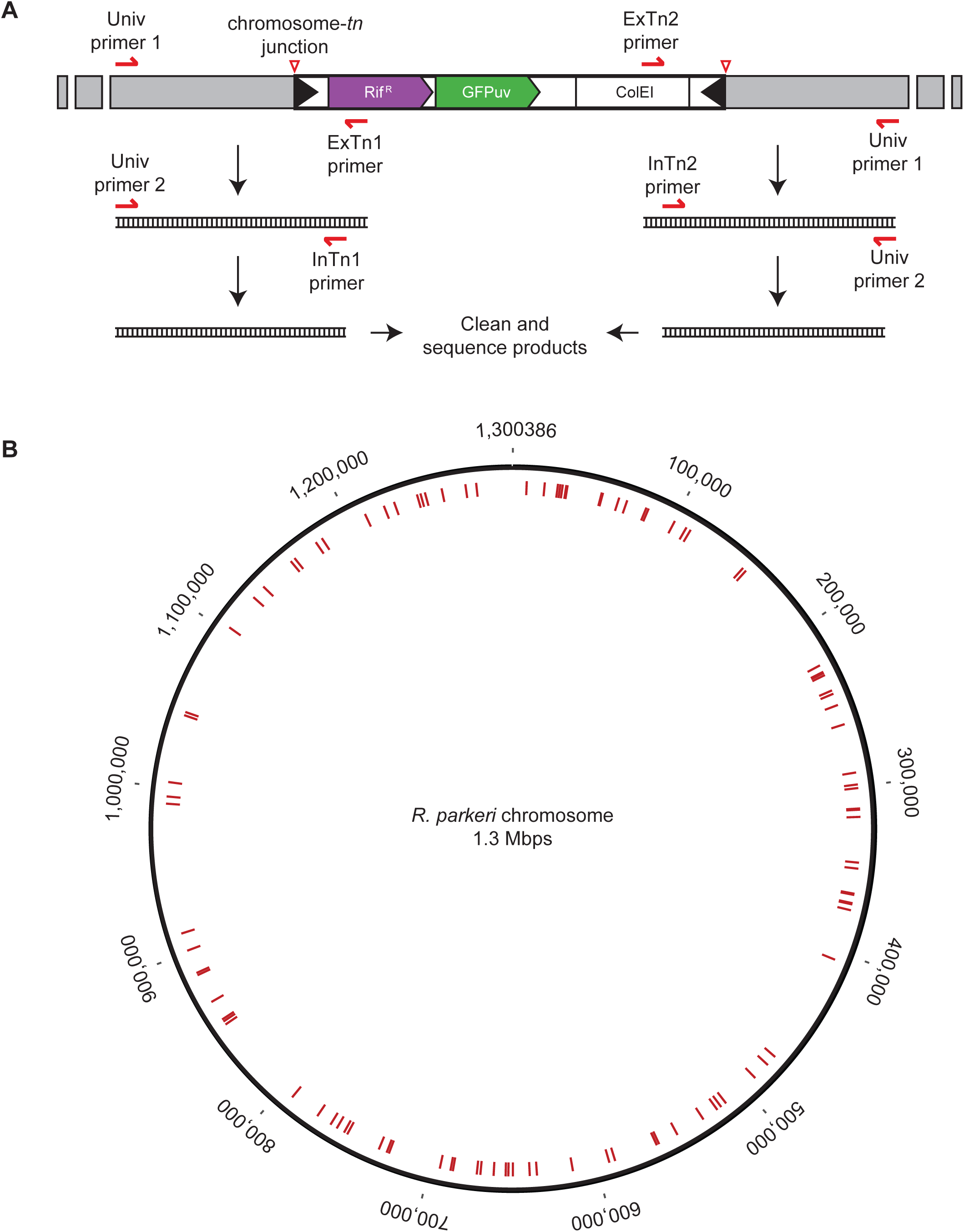
Mapping the transposon insertion sites. (A) Diagram showing the insertion of the transposon cassette into a chromosomal region (in grey). Primers specific to the transposon ends were paired with universal primers to amplify the chromosome-transposon junctions (red triangles), using semi-random nested PCR. Two nested PCR reactions were done to improve amplification of the chromosome-transposon junction directly from boiled bacteria. (B) *R. parkeri* chromosomal map showing all transposon insertion sites (see red lines) identified in this screen.

To identify genes involved in infection, we screened for transformants that showed altered plaque size and/or morphology (Fig 1B). We predicted that plaque size changes would result from defects at different stages of the rickettsial life cycle, including in intracellular growth, replication, motility, and/or spread. To screen for such mutants, pMW1650-electroporated bacteria were immediately used to setup plaque assays in the presence of rifampicin to select for transformants. Plaque size was monitored visually over the course of 3-9 days, and those displaying a small plaque (Sp) or big plaque (Bp) phenotype relative to their neighbors were selected for expansion. After independently repeating this process 7 times, 120 Sp mutant and 2 Bp mutants were selected for further analysis, as detailed below.

### Expansion and mapping of the transposon mutants

To expand the isolated transformants, plaques were picked and transferred to uninfected host cells to propagate each bacterial strain. Once the host cells were >75% infected, *Rickettsia* were purified using the rapid isolation procedure outlined above. Nine isolates did not grow in this expansion procedure, possibly due to a lack of live bacteria in the original plaque or poor isolation of the infected cells. The remaining transformants could be expanded, and for these the transposon insertion site was mapped using a semi-random nested PCR protocol. In short, the junctions between the transposon and the flanking genomic regions were amplified via two nested PCR reactions using transposon-specific and universal primers (Fig 1C). PCR products were sent directly for sequencing. Mapping of the transposon insertion sites to the *R. parkeri* chromosome (accession number CP003341) revealed no preference for specific regions (Fig 1D), similar to what was observed in *R. rickettsii* [18,19] and *R. prowazekii* [16,18]. Using this procedure, we identified the transposon insertion sites for 106 mutants. For 6 isolates the transposon insertion site could not be mapped, and the strains did not express GFP_uv_ (data not shown), suggesting these were spontaneous rifampicin-resistant strains. Of the 106 transposon mutations mapped, 81 were within the coding regions of 75 distinct genes and 25 were in intergenic regions (Table 1). Mutants of interest (Sp2, Sp34, and Sp19) were further purified and expanded for detailed analysis, as previously reported [14,20]. Our results indicate that transposon mutagenesis can be readily adapted for large-scale forward genetic screening to study gene function in *R. parkeri*.

**Table 1.** Transposon insertion sites in *R. parkeri*

| Name | Insertion site | Gene symbol | Gene product description |
| --- | --- | --- | --- |
| Sp1 | 101427 | MC1_00610 | Putative cytoplasmic protein |
| Sp2 | 112315 | MC1_00650 ** | Surface cell antigen 2 (Sca2) |
| Sp3 | 681322-681323 | MC1_03895 | Single-stranded-DNA-specific exonuclease RecJ |
| Sp6 | 365840 | MC1_02010 | Cytochrome c1, heme protein |
| Sp7 | 670632 | MC1_03810 | Folypolyglutamate synthase |
| Sp8 | 1047805-1047806 | Intergenic | n/a |
| Sp9 | 151196-151197 | MC1_00820 | VirB6 Type IV secretory pathway (rvhB6e) |
| Sp10 | 491813 | Intergenic | n/a |
| Sp11 | 1136147 | MC1_06660 | DNA polymerase I |
| Sp13 | 563189-563190 | MC1_03195 | RND efflux transporter |
| Sp14 | 518698-518699 | MC1_02960 | CTP synthetase |
| Sp15 | 520939 | MC1_02980 | Hypothetical protein |
| Sp17 | 1248850-1248851 | MC1_07220 | Transcriptional regulator |
| Sp18 | 70364 | MC1_00450 | Hypothetical protein |
| Sp19 | 654506-654507 | MC1_03740 | Antigenic heat-stable 120 kDa protein (Sca4) |
| Sp20 | 531536 | MC1_03025 ** | ampG protein |
| Sp21 | 20179 | Intergenic | n/a |
| Sp22 | 29609 | MC1_00175 | F0F1 ATP synthase subunit B |
| Sp23 | 474265 | MC1_02665 | Outer membrane assembly protein |
| Sp24 | 753916 | Intergenic | n/a |
| Sp25 | 728290 | MC1_04100 | Isopentenyl pyrophosphate isomerase |
| Sp26 | 33338 | MC1_00210 | Transcriptional regulator |
| Sp27 | 30722 | Intergenic | n/a |
| Sp28 | 301811 | MC1_01650 | Protease |
| Sp29 | 225255 | MC1_01180 | Acriflavin resistance protein D |
| Sp30 | 886852 | Intergenic | n/a |
| Sp31 | 262955-262956 | MC1_01410 | Hypothetical protein |
| Sp33 | 299510 | MC1_01640 | Putative toxin of toxin-antitoxin (TA) system |
| Sp34 | 888003 | MC1_05085 ** | Actin polymerization protein RickA |
| Sp35 | 589425 | MC1_03370 | Thiol:disulfide interchange protein dsbA |
| Sp36 | 230327 | MC1_01215 | Prolyl endopeptidase |
| Sp37 | 637085 | MC1_03670 | Hypothetical protein |
| Sp38 | 292360-292361 | MC1_01595 | S-adenosylmethionine:tRNA ribosyltransferase-isomerase |
| Sp39 | 912985 | MC1_05235 | Hypothetical protein (RARP-2) |
| Sp40 | 1279632 | Intergenic | n/a |
| Sp41 | 995818 | Intergenic | n/a |
| Sp42 | 372020 | MC1_02055 ** | GTP-binding protein LepA |
| Sp43 | 651603-651604 | MC1_03735 | ADP, ATP carrier protein |
| Sp44 | 868641 | MC1_04970 | HAD-superfamily hydrolase |
| Sp45 | 761156 | MC1_04295 | Microcin C7 resistance protein |
| Sp46 | 852817 | MC1_04870 | Methylated-DNA-protein-cysteine methyltransferase |
| Sp47 | 856486 | MC1_04920 | Hypothetical protein |
| Sp48 | 243782-243780 | MC1_01300 | DNA repair protein RecN |
| Sp49 | 687339 | MC1_03930 | Hypothetical protein |
| Sp50 | 1158028 | MC1_06730 | Hypothetical protein |
| Sp51 | 888088 | MC1_05085 ** | Actin polymerization protein RickA |
| Sp52 | 346470 | Intergenic | n/a |
| Sp53 | 793762 | MC1_04525 | Hypothetical protein |
| Sp54 | 1258878 | MC1_07285 | Hypothetical protein |
| Sp55 | 1245896 | MC1_07200 | tig Trigger factor |
| Sp56 | 1243741-1243742 | Intergenic | n/a |
| Sp57 | 1004249-1004250 | Intergenic | n/a |
| Sp58 | 1273429 | MC1_07360 | NAD(P) transhydrogenase subunit alpha |
| Sp59 | 1181227 | Intergenic | n/a |
| Sp60 | 1158028 | MC1_01115 | Hypothetical protein |
| Sp62 | 350030-350031 | MC1_01915 | Cytochrome c oxidase assembly protein |
| Sp63 | 109070 | MC1_00650 ** | Surface cell antigen (Sca2) |
| Sp64 | 314932 | MC1_01745 ** | Ankyrin repeat-containing protein (RARP-1) |
| Sp65 | 726967 | Intergenic | n/a |
| Sp66 | 615509-615510 | MC1_03545 | Hypothetical protein |
| Sp71 | 371351 | MC1_02055 ** | GTP-binding protein LepA |
| Sp72 | 83786 | MC1_00525 | Stage 0 sporulation protein J |
| Sp73 | 655844 | MC1_03745 | Putative transcriptional regulator |
| Sp74 | 991759-991760 | MC1_05745 | Hypothetical protein |
| Sp75 | 251100 | MC1_01335 | Ankyrin repeat-containing protein |
| Sp76 | 9674 | Intergenic | n/a |
| Sp78 | 672659 | Intergenic | n/a |
| Sp79 | 65481 | Intergenic | n/a |
| Sp80 | 1210788-1210789 | MC1_07040 | Outer membrane protein OmpA |
| Sp81 | 365135 | MC1_02000 | Cytochrome b |
| Sp82 | 549578-549579 | MC1_03115 | Cytochrome c oxidase polypeptide |
| Sp83 | 480829 | MC1_02715 | Hypothetical protein |
| Sp84 | 689140 | Intergenic | n/a |
| Sp85 | 514488 | Intergenic | n/a |
| Sp88 | 241435 | MC1_01295 | Thermostable carboxypeptidase |
| Sp90 | 1127301 | MC1_06610 | Hypothetical protein |
| Sp91 | 82796 | MC1_00515 | 16S rRNA methyltransferase GidB |
| Sp92 | 1229489-1229490 | MC1_07110 | 17 kDa surface antigen |
| Sp93 | 1223170 | MC1_07070 | Undecaprenyl-phosphate alpha-N-acetylglucosaminyltransferase |
| Sp94 | 774831 | Intergenic | n/a |
| Sp95 | 902617 | MC1_05150 | Patatin b1 |
| Sp96 | 561640-561641 | MC1_03180 | Hypothetical protein |
| Sp97 | 641129-641130 | MC1_03685 | miaA tRNA delta(2)-isopentenylpyrophosphate transferase |
| Sp98 | 34100-34101 | MC1_00220 | Putative methyltransferase |
| Sp99 | 1104365 | Intergenic | n/a |
| Sp100 | 375061 | Intergenic | n/a |
| Sp101 | 152889-152890 | Intergenic | n/a |
| Sp102 | 406474-406475 | MC1_02260 | DNA mismatch repair protein MutS |
| Sp103 | 662735 | MC1_03780 | Hypothetical protein |
| Sp104 | 1161553 | MC1_06745 | Hypothetical protein |
| Sp105 | 593543-593544 | MC1_03405 | Acylamino acid-releasing protein |
| Sp106 | 1045462-1045463 | MC1_06065 | Outer membrane protein OmpB |
| Sp107 | 531709 | MC1_03025 ** | ampG protein |
| Sp108 | 1177263 | MC1_06810 | F0F1 ATP synthase subunit beta |
| Sp109 | 54288-54289 | MC1_00370 ** | Chaperone ClpB |
| Sp111 | 854916 | Intergenic | n/a |
| Sp112 | 55657 | MC1_00370 ** | Chaperone ClpB |
| Sp113 | 319455 | MC1_01760 | Histidine kinase sensor protein |
| Sp114 | 765596 | MC1_04335 | Ribonuclease D |
| Sp115 | 229548 | Intergenic | n/a |
| Sp116 | 314408 | MC1_01745 ** | Ankyrin repeat-containing protein (RARP-1) |
| Sp117 | 733347 | MC1_04135 | Hypothetical protein |
| Sp118 | 695571 | Intergenic | n/a |
| Sp119 | 231259 | MC1_01235 | Prolyl endopeptidase |
| Sp120 | 27839 | MC1_00155 | Hypothetical protein |
| Bp2 | 756531 | MC1_04275 | Hypothetical protein |

Spontaneous rifampicin resistant mutants: Sp4-5, 69-70, 86, Bp1. Clones that did not expand: Sp16, 32, 61, 67-68, 77, 87, 89, 110. Mapping for Sp12 revealed two different insertion sites and was not included in the above list. ** Indicates more than one isolated transposon mutant/gene. n/a, not applicable.

## Discussion

A critical barrier to identifying and characterizing virulence factors in obligate intracellular bacterial pathogens has been the inability to easily manipulate their genomes. In this study, we sought to overcome this barrier and harness recent advances in rickettsial genetics to build a library of transposon mutants of the SFG *Rickettsia* species, *R. parkeri*. We streamlined previous protocols to more efficiently introduce a *mariner*-based transposon into the *R. parkeri* genome and isolated 106 independent transposon insertion mutations. Our study represents the first such transposon mutant library in this species, and the most extensive reported library in the rickettsiae.

In our study, we selected for mutants that showed an altered plaque size phenotype in infected host cell monolayers. Transposon mutations may cause a small plaque phenotype due to any number of defects, including: poor bacterial replication, reduced access to or survival within the cytosol, impaired cytosolic actin-based motility, and defective cell-to-cell spread. It was thus not surprising that we identified genes with a diverse set of predicted functions. Many genes with products predicted to perform bacterial-intrinsic functions (e.g. DNA replication) were identified and are expected to indirectly influence host-pathogen interactions through their role in bacterial growth and division. Other genes had more direct connections to the infectious life cycle and were further characterized in our recent studies to reveal their specific functions in intracellular infection [14,20]. For example, we previously described transposon mutations that disrupt the *rickA* (Sp34) and *sca2* (Sp2) genes and showed that these gene products are required for two independent phases of *R. parkeri* actin-based motility [20]. We also identified a transposon insertion (Sp19) in *sca4* gene and showed this encodes a secreted effector that promotes cell-to-cell spread [14].

Other genes mutated in this screen have been suggested to play critical roles during the infectious life cycle of other *Rickettsia* species but have yet to be characterized in *R. parkeri*. For example, we isolated transposon insertion mutants in the *ompA* (Sp80) and *ompB* (Sp106) genes, encoding the outer membrane proteins OmpA and OmpB. Work with SFG species *R. conorii* and *R. rickettsii* showed that OmpA and OmpB may regulate adhesion to and/or invasion of host cells [22-25]. However, some of this work relied on expression of these proteins in other bacterial species because mutants were not available. Interestingly, targeted knockout of *ompA* in *R. rickettsii* via an LtrA group II intron retrohoming system revealed no clear requirement for OmpA in invasion [26], suggesting it alone is not necessary. This highlights the importance of studying loss-of-function mutants to reveal gene function. The fact that *ompA* and *ompB* mutants exhibit a small plaque phenotype suggest additional functions of these proteins, putative indirect effects of the truncated products, or simply a need for efficient invasion of neighboring cells after host cells lyse during plaque development. Future work on these mutants should help reveal the relative importance of these proteins during invasion and/or other stages of the *R. parkeri* life cycle.

Our screen also revealed genes for which no specific role has been ascribed during the infectious life cycle, although the sequence of their protein products suggests a role in interaction with host cells. These proteins include some with eukaryotic-like motifs such as ankyrin repeats, which often mediate protein-protein interactions [27], and are a common motif in secreted bacterial effector proteins or virulence factors that target host pathways [27,28]. In particular, mutations in genes encoding *R. parkeri* orthologs of RARP-1 and RARP-2 from *R. typhi* (accession numbers MC1_01745 and MC1_05235, respectively) were identified in our screen (Sp64, Sp116, and Sp39). Work in *R. typhi* has revealed that RARP-1 and RARP-2 are secreted into the host cell, but their precise functions remain unknown [29,30].

Another mystery in rickettsial biology relates to the functional importance of the putative type IV secretion system (T4SS) encoded in their genomes [31], which in other species is involved in translocating DNA, nucleoproteins, and effector proteins into host cells [32]. Strikingly, the *Rickettsia* T4SS has an unusual expansion of the VirB6-like genes (i.e. Rickettsiales vir homolog, *rvhB6*), which are predicted to encode inner membrane protein components at the base of the T4SS [30,31,33]. Interestingly, we isolated a strain with a transposon insertion mutation in the fifth VirB6-like gene, *rvhB6e* (Sp9). This mutant will prove useful to explore the function of the T4SS in rickettsial infection.

Finally, we identified 20 strains, each carrying a transposon insertion disrupting a gene encoding a hypothetical protein. One of these caused a big plaque phenotype, suggesting enhanced growth or cell-to-cell spread. Further study of these mutants has the potential to reveal the function of these uncharacterized gene productions during rickettsial infection.

Overall, our mutant collection provides an important resource that can be used to uncover key bacterial players that regulate rickettsial interactions with their host cells. By streamlining the ease of mutant isolation as described here, investigators can begin expanding the available toolkit to generate more *Rickettsia* mutants. This will also allow for more direct analysis of gene function in the rickettsiae without the reliance on introducing genes into heterologous organisms. This forward genetics approach has the potential to reveal new insights into rickettsial biology and pathogenesis; however, limitations remain. For example, because the rickettsiae are obligate intracellular pathogens, screens such as these are unlikely to reveal genes that are essential for invasion or intracellular growth. Therefore, we cannot necessarily assess the relative importance of genes not identified in forward genetic screens. Additionally, the reported protocol still has limits with regard to electroporation efficiency and recovery on host cells, which makes it harder to adapt for large-scale mutagenesis work. Further advancements in rickettsial genetic methods will be necessary to complement our work and more effectively probe the molecular mechanisms of all genes whose products may control critical host-pathogen interactions.

## Acknowledgments

We thank David Wood and Chris Paddock for plasmids and strains and Shawna Reed for technical help. We also thank the staff of UC Berkeley’s Cell Culture Facility and DNA Sequencing Facility for critical experimental support. R.L.L. was supported by a Helen Hay Whitney Foundation postdoctoral fellowship and NIH/NIGMS grant K99 GM115765. M.D.W. is supported by NIH/NIAID grant R01 AI109044.

## References

1 Gillespie JJ, Williams K, Shukla M, Snyder EE, Nordberg EK, Ceraul SM, et al. *Rickettsia* phylogenomics: unwinding the intricacies of obligate intracellular life. PLoS ONE. 2008;3: e2018. doi:10.1371/journal.pone.0002018

2 Walker DH, Ismail N. Emerging and re-emerging rickettsioses: endothelial cell infection and early disease events. Nat Rev Microbiol. 2008;6: 375–386. doi:10.1038/nrmicro1866

3 Paddock CD, Sumner JW, Comer JA, Zaki SR, Goldsmith CS, Goddard J, et al. *Rickettsia parkeri*: a newly recognized cause of spotted fever rickettsiosis in the United States. Clin Infect Dis. 2004;38: 805–811. doi:10.1086/381894

4 Paddock CD, Finley RW, Wright CS, Robinson HN, Schrodt BJ, Lane CC, et al. *Rickettsia parkeri* rickettsiosis and its clinical distinction from Rocky Mountain spotted fever. Clin Infect Dis. 2008;47: 1188–1196. doi:10.1086/592254

5 Grasperge BJ, Reif KE, Morgan TD, Sunyakumthorn P, Bynog J, Paddock CD, et al. Susceptibility of inbred mice to *Rickettsia parkeri*. Infect Immun. 2012;80: 1846–1852. doi:10.1128/IAI.00109-12

6 Banajee KH, Embers ME, Langohr IM, Doyle LA, Hasenkampf NR, Macaluso KR. *Amblyomma maculatum* feeding augments *Rickettsia parkeri* infection in a rhesus macaque model: A pilot study. PLoS ONE. 2015;10: e0135175. doi:10.1371/journal.pone.0135175

7 Ray K, Marteyn B, Sansonetti PJ, Tang CM. Life on the inside: the intracellular lifestyle of cytosolic bacteria. Nat Rev Microbiol. 2009;7: 333–340. doi:10.1038/nrmicro2112

8 Lamason RL, Welch MD. Actin-based motility and cell-to-cell spread of bacterial pathogens. Curr Opin Microbiol. 2017;35: 48–57. doi:10.1016/j.mib.2016.11.007

9 Reed SCO, Serio AW, Welch MD. *Rickettsia parkeri* invasion of diverse host cells involves an Arp2/3 complex, WAVE complex and Rho-family GTPase-dependent pathway. Cell Microbiol. 2012;14: 529–545. doi:10.1111/j.1462- 5822.2011.01739.x

10 Welch MD, Reed SCO, Haglund CM. Establishing intracellular infection: escape from the phagosome and intracellular colonization (rickettsiaceae). In: Palmer GH Azad AF, editors. Washington DC: Intracellular Pathogens II: Rickettsiales; 2012. pp. 154–174. doi:10.1128/9781555817336.ch5

11 Heinzen RA, Hayes SF, Peacock MG, Hackstadt T. Directional actin polymerization associated with spotted fever group *Rickettsia* infection of Vero cells. Infect Immun. 1993;61: 1926–1935.

12 Serio AW, Jeng RL, Haglund CM, Reed SC, Welch MD. Defining a core set of actin cytoskeletal proteins critical for actin-based motility of *Rickettsia*. Cell Host Microbe. 2010;7: 388–398. doi:10.1016/j.chom.2010.04.008

13 Choe JE, Welch MD. Actin-based motility of bacterial pathogens: mechanistic diversity and its impact on virulence. Carbonetti N, editor. Pathog Dis. 2016;74: pftw099. doi:10.1093/femspd/ftw099

14 Lamason RL, Bastounis E, Kafai NM, Serrano R, Del Álamo JC, Theriot JA, et al. *Rickettsia* Sca4 reduces vinculin-mediated intercellular tension to promote spread. Cell. 2016;167: 670–683.e10. doi:10.1016/j.cell.2016.09.023

15 McClure EE, Chávez ASO, Shaw DK, Carlyon JA, Ganta RR, Noh SM, et al. Engineering of obligate intracellular bacteria: progress, challenges and paradigms. Nat Rev Microbiol. 2017;15: 544–558. doi:10.1038/nrmicro.2017.59

16 Liu Z-M, Tucker AM, Driskell LO, Wood DO. Mariner-based transposon mutagenesis of *Rickettsia prowazekii*. Appl Environ Microbiol. 2007;73: 6644–6649. doi:10.1128/AEM.01727-07

17 Qin A, Tucker AM, Hines A, Wood DO. Transposon mutagenesis of the obligate intracellular pathogen *Rickettsia prowazekii*. Appl Environ Microbiol. 2004;70: 2816–2822.

18 Clark TR, Lackey AM, Kleba B, Driskell LO, Lutter EI, Martens C, et al. Transformation frequency of a mariner-based transposon in *Rickettsia rickettsii*. 2011;193: 4993–4995. doi:10.1128/JB.05279-11

19 Kleba B, Clark TR, Lutter EI, Ellison DW, Hackstadt T. Disruption of the *Rickettsia rickettsii* Sca2 autotransporter inhibits actin-based motility. Infect Immun. 2010;78: 2240–2247. doi:10.1128/IAI.00100-10

20 Reed SCO, Lamason RL, Risca VI, Abernathy E, Welch MD. *Rickettsia* actin-based motility occurs in distinct phases mediated by different actin nucleators. Curr Biol. 2014;24: 98–103. doi:10.1016/j.cub.2013.11.025

21 Welch MD, Reed SCO, Lamason RL, Serio AW. Expression of an epitope-tagged virulence protein in *Rickettsia parkeri* using transposon insertion. PLoS ONE. 2012;7: e37310. doi:10.1371/journal.pone.0037310

22 Uchiyama T. Adherence to and invasion of Vero cells by recombinant *Escherichia coli* expressing the outer membrane protein rOmpB of *Rickettsia japonica*. Ann N Y Acad Sci. 2003;990: 585–590.

23 Li H, Walker DH. rOmpA is a critical protein for the adhesion of *Rickettsia rickettsii* to host cells. Microb Pathog. 1998;24: 289–298. doi:10.1006/mpat.1997.0197

24 Hillman RD, Baktash YM, Martinez JJ. OmpA-mediated rickettsial adherence to and invasion of human endothelial cells is dependent upon interaction with α2β1 integrin. Cell Microbiol. 2013;15: 727–741. doi:10.1111/cmi.12068

25 Chan YGY, Cardwell MM, Hermanas TM, Uchiyama T, Martinez JJ. *Rickettsial* outer-membrane protein B (rOmpB) mediates bacterial invasion through Ku70 in an actin, c-Cbl, clathrin and caveolin 2-dependent manner. Cell Microbiol. 2009;11: 629–644. doi:10.1111/j.1462-5822.2008.01279.x

26 Noriea NF, Clark TR, Hackstadt T. Targeted knockout of the *Rickettsia rickettsii* OmpA surface antigen does not diminish virulence in a mammalian model system. Bio. 2015;6. doi:10.1128/mBio.00323-15

27 Al-Khodor S, Price CT, Kalia A, Abu Kwaik Y. Functional diversity of ankyrin repeats in microbial proteins. Trends Microbiol. 2010;18: 132–139. doi:10.1016/j.tim.2009.11.004

28 Pan X, Lührmann A, Satoh A, Laskowski-Arce MA, Roy CR. Ankyrin repeat proteins comprise a diverse family of bacterial type IV effectors. Science. 2008;320: 1651–1654. doi:10.1126/science.1158160

29 Kaur SJ, Rahman MS, Ammerman NC, Beier-Sexton M, Ceraul SM, Gillespie JJ, et al. TolC-dependent secretion of an ankyrin repeat-containing protein of *Rickettsia typhi*. J Bacteriol. 2012;194: 4920–4932. doi:10.1128/JB.00793-12

30 Gillespie JJ, Kaur SJ, Rahman MS, Rennoll-Bankert K, Sears KT, Beier-Sexton M, et al. Secretome of obligate intracellular *Rickettsia*. FEMS Microbiol Rev. 2015;39: 47–80. doi:10.1111/1574-6976.12084

31 Gillespie JJ, Ammerman NC, Dreher-Lesnick SM, Rahman MS, Worley MJ, Setubal JC, et al. An anomalous type IV secretion system in *Rickettsia* is evolutionarily conserved. PLoS ONE. 2009;4: e4833. doi:10.1371/journal.pone.0004833

32 Grohmann E, Christie PJ, Waksman G, Backert S. Type IV secretion in Gram-negative and Gram-positive bacteria. Mol Microbiol. 2018;107: 455–471. doi:10.1111/mmi.13896

33 Gillespie JJ, Phan IQH, Driscoll TP, Guillotte ML, Lehman SS, Rennoll-Bankert KE, et al. The *Rickettsia* type IV secretion system: unrealized complexity mired by gene family expansion. Mobley H, editor. Pathog Dis. 2016;74: pftw058. doi:10.1093/femspd/ftw058

